# SoluProtMut: Siamese Deep Learning for Predicting Solubility Effects of Protein Mutations with Experimental Validation

**DOI:** 10.1101/2025.09.26.676459

**Authors:** Jan Velecky, Hana Faldynova, Pedro Hermosilla, Katerina Slanska, Nela Sendlerova, Mark Doerr, Sara Egersdorfova, Uwe Bornscheuer, Jiří Damborsky, Zbynek Prokop, Stanislav Mazurenko

**Author notes:** joint first authors (Jan Velecky, Hana Faldynova). corresponding authors (Z. Prokop) and (S. Mazurenko).

## Abstract

Protein solubility is an attractive engineering target because it is a critical property influencing the scalability of protein production and the success of therapeutic proteins in biomedical applications. However, predicting *in silico* solubility changes upon mutation is challenging due to data heterogeneity and protein bias. Here, we explore how different sources of solubility data can be used for machine learning and present SoluProtMut, a Siamese deep geometric neural network trained to predict the impact of mutations on protein solubility. Our final model was trained exclusively on deep mutational scanning data derived from recombinant protein expression assays in bacterial and yeast systems. We compare our model with five established solubility prediction methods. On an independent set of 44 proteins, the model achieves the highest informedness among the methods compared, with the largest margin for multipoint mutations (a ternary informedness of 26.5%, 8.8 percentage points above the next-best method). Our findings also reaffirm that the scarcity of solubility data continues to hamper progress in this field. To address this limitation, we experimentally quantified solubility changes for 277 single-point and multipoint mutants of haloalkane dehalogenase. Complemented by recent deep-mutational-scanning data on myoglobin, we employed both datasets for external validation. We identify protein-level generalization, rather than model capacity or training-set size, as the main limiting factor in the domain. Integrating high-throughput assays with deep learning nonetheless offers a route to improving the accuracy and scope of solubility prediction.

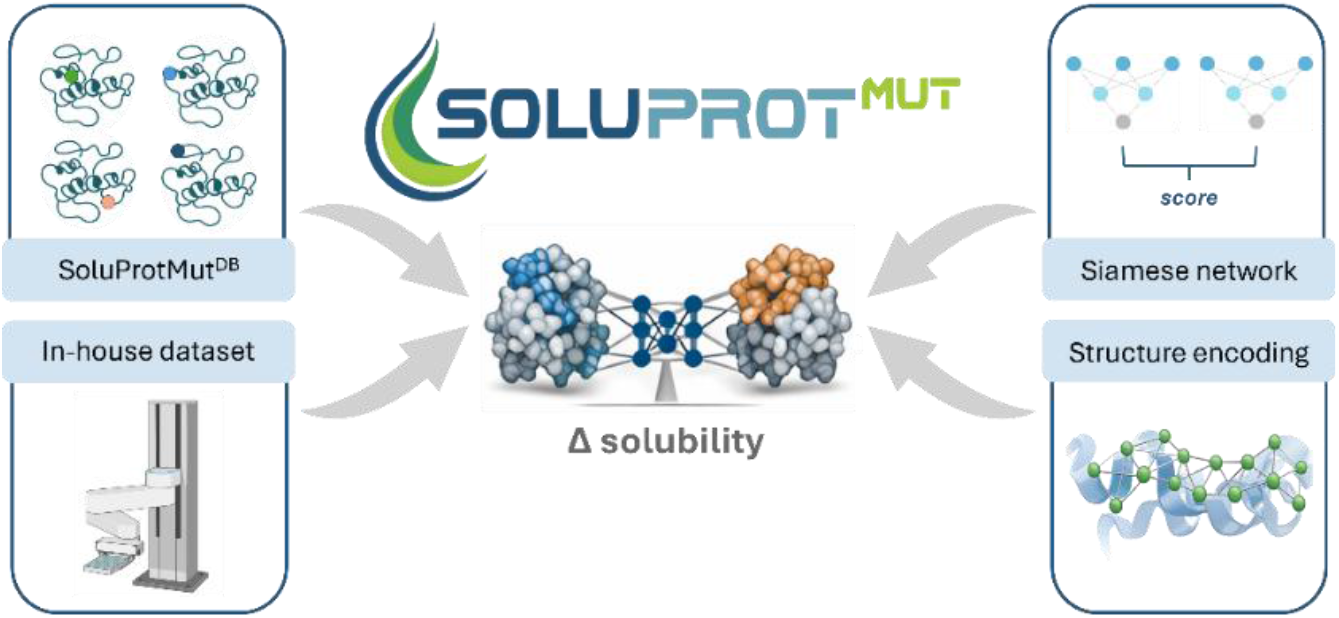

## Introduction

Protein solubility is a fundamental property of globular proteins with broad implications in both biotechnology and medicine. It is the maximum concentration at which a protein remains freely dissolved in solution under given conditions (Kramer *et al*., 2012). Low solubility often limits the yield of recombinant protein production, impacting the economy of industrial applications. It may also complicate the administration of protein biopharmaceuticals in sufficient dosing (Shire *et al*., 2004; Rosace *et al*., 2023). As such, solubility has become an important target in protein design.

Traditionally, solubility of mutant designs is assessed using classical biochemical assays, such as Native-PAGE or SDS-PAGE. More recently, high-throughput approaches emerged, particularly those involving deep mutational scanning (DMS), which allows a rapid and systematic screening of thousands of protein variants. These approaches often employ green fluorescent protein (GFP) with a fluorescence signal as a proxy for solubility. One of these approaches utilizes the complementation of GFP fragments (Cabantous, Terwilliger, *et al*., 2005; Cabantous, Pédelacq, *et al*., 2005), where a small non-fluorescent GFP fragment is fused to the protein of interest, and only upon expression and correct folding, the fluorescence is detected. Despite these advances, experimental approaches remain labour-intensive or costly, motivating the development of accurate *in silico* predictors.

Early efforts to predict protein solubility began with the Wilkinson–Harrison model, which used a logistic regression to estimate the likelihood of soluble expression in *Escherichia coli* (*E. coli*) (Wilkinson and Harrison, 1991). Later, this model was revised to consider only turn-forming and charged residues (Davis *et al*., 1999). Afterwards, the development of solubility prediction methods split into two branches: (a) absolute predictors, which classify the protein as soluble or insoluble, and (b) mutational predictors, which assess the impact of specific mutations. Several absolute predictors are widely used today, such as SOLpro (Magnan *et al*., 2009), PROSO II (Smialowski *et al*., 2012) or SoluProt (Hon *et al*., 2021), to name a few. These methods are typically validated for expression in *E. coli*, offering a strategy to assess the viability of heterologous expression.

In contrast, mutational predictors are less common. Notable examples include OptSolMut (Tian *et al*., 2010), the first computational model and the first dataset for this task; and PON-Sol2 (Yang *et al*., 2021), which significantly expanded the training data by incorporating the available DMS data. A middle ground between absolute and mutational predictors is CamSol (Sormanni *et al*., 2015), which provides solubility profiles to guide site-specific design.

A related class of computational approaches focuses on protein design. For example, PROSS (Goldenzweig *et al*., 2016) combines evolutionary information with atomistic modelling to propose multipoint variants with improved stability, expression, and solubility. Unlike mutational solubility predictors, which estimate the effect of a specified amino acid substitution, such design-oriented frameworks directly generate candidate variants.

The best reported blind-test performance for mutational solubility prediction remains modest, with a normalized correct prediction ratio of 0.545 reported for PON-Sol2, highlighting substantial room for improvement (Yang *et al*., 2021; Wee *et al*., 2024). One of the major challenges is the training data heterogeneity, as annotations for protein solubility come from a wide selection of experimental assays – from low-throughput experiments for many proteins, often with precise quantifications, to high-throughput enrichment ratios collected for only a few proteins (Velecký *et al*., 2022).

Here, we introduce SoluProtMut, a model for predicting the impact of mutations on protein solubility. With SoluProtMut, we delved into various solubility datasets to explore their potential for machine learning. Our final model was trained exclusively on the DMS-derived data. SoluProtMut builds upon the IEConv architecture, originally developed for protein representation learning (Hermosilla *et al*., 2020). To adapt the model for mutational prediction, we employed a Siamese architecture (Chicco, 2021), a strategy previously used in a neighbouring field of mutational stability prediction (Zhou *et al*., 2023; Zhang *et al*., 2025). Our Siamese design enforces anti-symmetry (Sanavia *et al*., 2020) through a differential connection, diverging from the prior works that rely on regularization terms in the loss function. Our model outperformed the previous state of the art and displayed higher precision in identifying desolubilizing mutations, which is critical for avoiding detrimental designs in protein engineering. In addition, we assembled two new datasets for independent validation: myoglobin from the previously published DMS data (Küng et al., 2025), and LinB haloalkane dehalogenase, a new split-GFP-based dataset collected in-house. Our study revealed an evaluation dependence on a particular protein, highlighting the need for broader mutational solubility data covering unrepresented proteins.

## Methods

### Primary data source

The mutational solubility data used in this study comprise protein sequences, specific mutations, and their experimentally observed effects on solubility. Each data point thus represents a pair of protein variants (original and mutant) measured under equivalent experimental conditions, and importantly, within a single study. We acquired such data from SoluProtMut^DB^ (Velecký *et al*., 2022), a curated mutational solubility database. The exported data (dump date: May 27, 2024) contains 32 992 data points across 103 proteins, 72 of which have a structure assigned from the Protein Data Bank. Each data point is labelled with one of the discrete solubility effect classes: highly desolubilizing (−−), desolubilizing (−), neutral (N), solubilizing (+), and highly solubilizing (++).

It is worth noting that the database is class-imbalanced, with the desolubilizing cases substantially overrepresented over the solubilizing ones in a ratio of 7:2 (**Table S3**). Additionally, the class distribution among protein variants is typically highly skewed towards one direction. For instance, only desolubilizing and only solubilizing mutations are reported for 6 and 13 proteins, respectively.

Over 97 % of the data (32 081 data points) originates from deep mutational scanning (DMS) experiments, which combine the generation of large mutational libraries, high-throughput population filtering, and deep sequencing to explore the mutational landscape. Three proteins were subjects of DMS campaigns (Klesmith *et al*., 2017; Wrenbeck *et al*., 2019): levoglucosan kinase (LGK) from *Lipomyces starkeyi*, pyrrolidine ketide synthase (PKS) from *Atropa belladonna,* and beta-lactamase (TEM) from *E. coli*. Each protein was assessed in two distinct assays, resulting in duplicated variants. It is important to note that high-throughput assay scores serve only as proxies for actual solubility. For instance, they may as well correlate highly with thermodynamic stability (Klesmith et al., 2017).

### Training and test data. \

We divided the SoluProtMut^DB^ data into DMS and non-DMS sets, based on whether DMS was used in the experimental assay. The DMS data are well-suited for deep learning thanks to their abundance and relatively unbiased coverage of the mutational landscape. This should mitigate overfitting to biases often observed in low-throughput methods: specific protein outcomes (e.g., highly soluble or insoluble proteins), particular amino acid substitutions (e.g., data from alanine scanning), or specific sequence positions (e.g., data from site-directed mutagenesis). In contrast, the non-DMS data are characterized by greater protein diversity, providing a more representative benchmark for assessing the generalization of a model across unseen proteins. Thus, the DMS data were used primarily for the main training, while the non-DMS data were intended for either fine-tuning or final model evaluation. The data splits and the overall study design are illustrated in **Figure 1E**.

**Figure 1:**
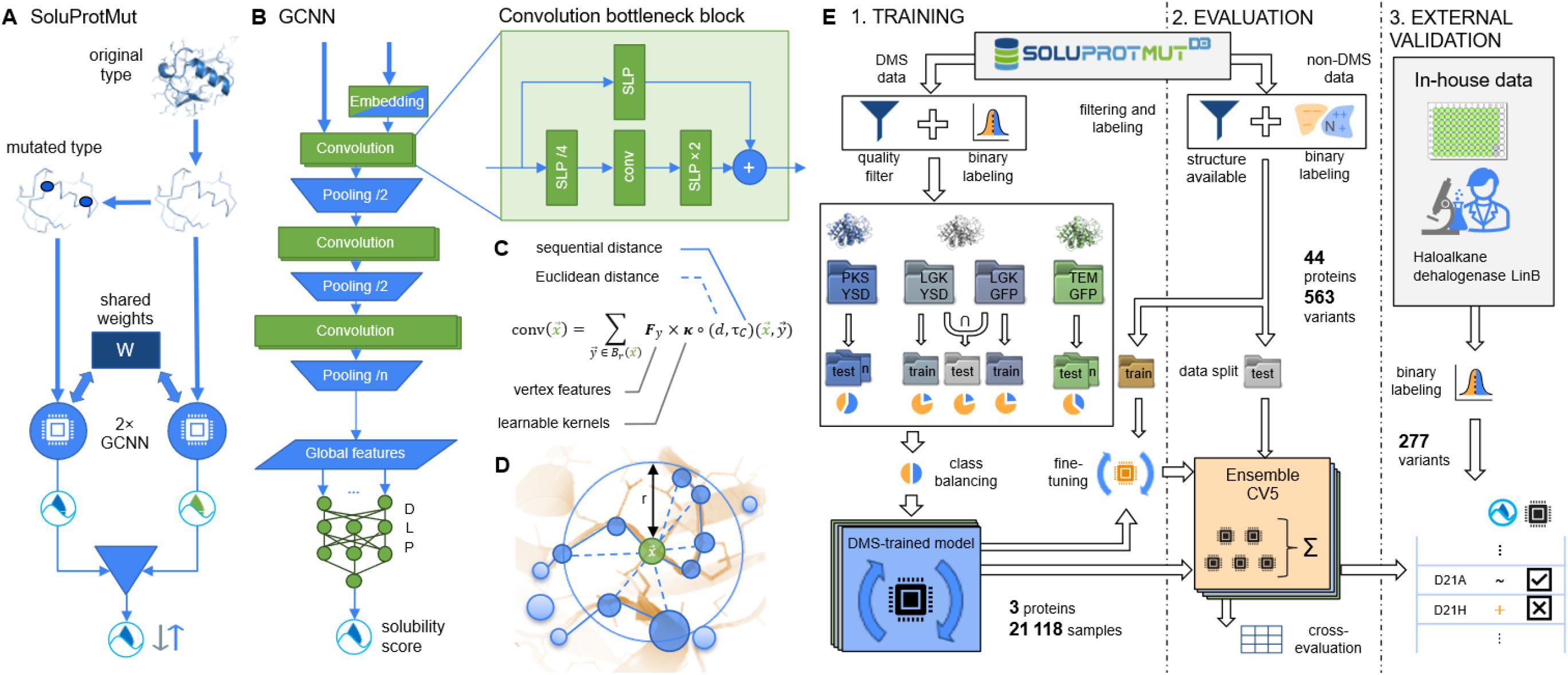
Design of this study. A: High-level architecture. Siamese network. Our change predictor consists of two identical base networks (GCNN) with shared weights, one fed with a wild-type and the other with a mutant. The classification is based on the difference between the predicted solubility scores. **B: Low-level architecture**. The base network (GCNN) consists of an encoding layer, three rounds of convolution and pooling layers, and a double-layer perceptron that implements the final classification layer and produces a solubility score. Each convolution layer is doubled and utilizes a bottleneck connection. The green nodes are learnable. **C: Convolution formula** for the convolution step over the vertices closer than r Å (Br) from the vertex 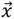. Kernels are learnable and parametrized by both sequential and Euclidean distances between the vertices**. D: Protein representation.** The structures within the network are represented as linear point clouds, enabling on-the-fly construction of a multigraph at each convolution step. On-the-fly edges (dashed) connect close vertices in Euclidean space. **E: Data processing.** We conducted the study in three phases: training, evaluation, and independent validation. We drew data from SoluProtMut^DB^ after filtering and binarizing the classes. We divided these data into deep mutational scanning data (DMS) and non-DMS data. The DMS data come from three proteins (PKS, LGK, and TEM) and two different assays (YSD and GFP), resulting in four experimental setups. We used these data to create several data splits and train multiple models in 5-fold cross-validation, yielding a 5-ensemble predictor. We created a single test set for LGK, comprising variants that consistently affect solubility in both assays. We weight-balanced each data split to counteract a higher abundance of the desolubilizing examples. The non-DMS data were used either to fine-tune the models or to evaluate them. Finally, we collected solubility data for 277 haloalkane dehalogenase variants in a wet lab, using a split-GFP medium-throughput assay, to independently validate the resulting model.

### DMS data preprocessing

To ensure the quality and consistency of the training data, we excluded data points originating from the Tat-export assay, following the authors’ observation about a high false detection rate of solubilizing variants (Klesmith *et al*., 2017). This only marginally reduced the number of available variants due to the overlap in the coverage by a second assay for each protein. After this exclusion, LGK was the only protein with duplicated variants labelled by two different assays – GFP and yeast surface display (YSD). The overview of the used DMS data is in **Table 1**. Next, we binarized the effects into two classes: negative (desolubilizing) and positive (non-desolubilizing), using a threshold of −10% on the assay-normalized fitness score, established in previous works (Klesmith *et al*., 2017; Velecký *et al*., 2022). This helped alleviate the imbalance between the number of solubilizing and desolubilizing mutations.

**Table 1:** Overview of deep mutational scanning experiments. These four DMS experiments provide training data of three proteins for the SoluProtMut predictor. The data comes from two different high-throughput assays (and also, host organisms), with LGK screened in both. The number of mutations roughly corresponds to the size of a respective protein with coverage from 66 % (PKS) up to 83 % (TEM). PKS stands out from the other two proteins with a high occurrence of rather solubilizing mutations.

| **Protein** | **Pyrrolidine ketide synthase (PKS)** | **Levoglucosan kinase (LGK)** |  | **β-lactamase (TEM)** |
|---|---|---|---|---|
| **Species** | *Atropa belladonna* | *Lipomyces starkeyi* |  | *Escherichia coli* |
| **Protein size (AAs)** | 388 | 439 |  | 263 |
| **Expression host** | *Escherichia coli* |  | *Saccharomyces cerevisiae* |  |
| **Solubility assay** | Fluorescence-activated cell sorting (GFP) |  | Yeast surface display (YSD) |  |
| **Mutation # (coverage)** | 4840 (66 %) | 6485 (78 %) | 6298 (75 %) | 4129 (83 %) |
| **Of which solubilizing** | 29.3 % | 6.6 % | 4.9 % | 10.3 % |
| **Modification** | GFP-tag |  | Epitope tag |  |

We created stratified training and test subsets, with 12 % of the data reserved for testing, for three assays: PKS, TEM and LGK-GFP. Furthermore, we created a curated LGK dataset by sampling data points that exhibit consistent solubility effects across both assays (YSD and GFP). The remaining data points were used for training. Additionally, we created a split for the entire DMS dataset by combining the respective splits of each DMS protein. Since PKS has no experimentally determined structure, we employed a homology model generated using I-TASSER with default parameters (Yang *et al*., 2015). To counteract the class imbalance, we applied class weights to the created subsets, as training on unweighted data often led the model to predict the majority class. For the combined DMS set, we further rebalanced the weights protein-wise to ensure equal contribution from each protein to training.

### Non-DMS data preprocessing

As our chosen network architecture learns on structures, we first filtered out all examples without assigned structures or with amino acids other than wild-type at the mutated positions. This resulted in 563 data points of 44 proteins in the dataset. We then split the data into training and test subsets as follows. Proteins observing mixed effects (both solubilizing and desolubilizing) for different mutations were kept for fine-tuning. Consequently, the holdout test set comprised the remaining proteins with consistent or neutral effects, thereby also including all single-data-point proteins. The split was designed to prevent the model from learning protein-specific biases.

To simplify the learning task, we binned non-DMS data into the positive (N, +, ++) and negative (−−, −) classes. Unlike the DMS data, no class- or protein-wise weighting was applied here due to the highly varying number of recorded examples per protein (from one to dozens) and many proteins featuring only one direction of effect for all their variants.

### Network design

SoluProtMut is a deep learning model designed to predict qualitative changes between two protein variants (wild-type and mutant). We adopted a Siamese architecture, which consists of a pair of identical base-model instances that share weights (**Figure 1A**). Each instance processes one of the two input structures and produces a score *S* ∈ [0,1]. The rescaled difference between the two scores, 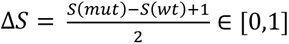, is a basis for a binary prediction. Mutations with ΔS below and above 0.5 are classified as desolubilizing and non-desolubilizing, respectively. Because swapping the two inputs returns 1 − ΔS, the predictor is anti-symmetric by construction rather than by penalty. To improve robustness, the final predictor is implemented as an ensemble of five base model pairs, trained independently in correspondence to the five cross-validation folds (CV5, see training procedure). The final output is determined by simple average voting, i.e., producing more stable output by averaging the predictions of the individual models.

The underlying regressor of SoluProtMut is a geometric neural network tailored for learning from protein structures. It is a graph-based convolutional neural network (GCNN) that leverages insights from the 3D organization of proteins and is based on the IEConv architecture (Hermosilla *et al*., 2020), initially developed for structural classification tasks such as EC number or fold classification. The network operates in three logical stages (**Figure 1B**): (1) processing and encoding of the input protein structure, (2) hierarchical convolution, and (3) final regression.

First, a structure representation is derived from the backbone coordinates provided in the input PDB file (**Figure 1D**). The coordinates are normalized so that the bounding box of the point cloud is centred at the origin of the coordinate system. The model uses wild-type backbones (C-α atoms) and mutant sequences as inputs. This approach eliminates the need for mutant structures and makes the trained model robust to side-chain conformational changes or optimizations.

The model input can be understood either as a linear vertex-weighted 3D graph or, equivalently, as a linear (ordered) point cloud. In both, each vertex or point initially corresponds to an amino acid, embedded in 3D space by its C-α atom. The vertex weights encode residue types, while the edges represent peptide bonds between residues. Each residue-type information is encoded using an 8-dimensional VHSE physicochemical descriptor (Mei *et al*., 2005). Thanks to the provided chemically meaningful features, the need to learn basic amino-acid properties from scratch is eliminated – a key distinction from the original IEConv design.

Second, the structure representation undergoes rounds of convolution and pooling. Each round consists of two convolutional layers followed by a pooling layer. A special convolution operation (described further) is performed in the 3D neighbourhood of each vertex, which initially represents an amino acid and later becomes a meta-residue after pooling. Pooling halves the number of vertices via average feature pooling, with new coordinates computed as the midpoint between each pair of vertices. The neighbourhood expands progressively at each stage, enabling the network to capture both local and global structural patterns. The last average pooling operation operates over all vertices, producing a global feature vector.

Each representation in convolution is treated as an edge-weighted multigraph, generated on-the-fly from the point cloud representation (**Figure 1D**). The edges are generated between all vertices within Euclidean distance of r, progressively increasing from 8 to 16 Å with every convolution round. Importantly, the convolution kernel is trainable and parameterized by the linear distance (i.e., sequential) and the Euclidean distance between vertices (**Figure 1C**). This allows the network to distinguish between spatially close residues adjacent or distant in the primary sequence. The convolution layers are organized into residual bottleneck blocks surrounding the actual convolution operation, inspired by ResNet (He *et al*., 2016). This enables efficient propagation of local features to subsequent layers (**Figure 1B**). Since the convolution operates on these graphs, it is also rotationally and translationally invariant to the input structure.

As the last step, the final score is produced by a double-layer perceptron (DLP) that operates on global features. This resulting regression value implicitly represents a solubility of the input structure, emerging from the differential training objective.

To improve generalization and training stability, batch normalization layers are applied after each linear layer. The entire network has 2.6 million trainable parameters. A summary of key modifications implemented over the original IEConv study is provided in **Table S2**.

### Training procedure

The initial training of the models was conducted in a five-fold cross-validation (CV5) setup, resulting in five independently trained models. The models were trained using a fixed learning-rate scheduler with three stages. We used minibatch stochastic gradient descent with binary cross-entropy as the loss function, comparing the predicted solubility effect against the true class. For each fold, the model with the highest validation accuracy was retained for evaluation. To regularize training, we applied weight decay (5e-4). We augmented the training data by perturbing the C-α coordinates of each input pair with Gaussian noise with zero mean and a standard deviation of 0.05 Å per coordinate. Crucially, identical perturbations were applied to both the wild-type and mutant structures to preserve the relative geometry.

We conducted fine-tuning experiments in two ways. In the full-model retraining, we started from a model trained on any two of the three DMS proteins and fine-tuned the entire network on the third, previously unseen protein. In the partial-model fine-tuning using only the non-DMS training subset, all layers were frozen except for the final regression module.

### Independent data collection

We used haloalkane dehalogenase LinB (UniProt: D4Z2G1) as a target to collect data on mutational effects on solubility using the split-GFP assay (Waldo *et al*., 1999). In this assay, adjusted for solubility, GFP is split into two non-fluorescent fragments: GFP_1-10_ (detector) and GFP_11_ (fusion tag). The tag is expressed in a fusion with a protein of interest, while the detector is produced and purified separately. Fluorescence is measured after co-incubation of the two fragments. This assay reports relative accumulation of soluble protein through GFP fragment complementation and was used here to quantify mutational effects on soluble expression rather than absolute protein yields.

A LinB substitution library with a GFP_11_ linker was constructed via error-prone PCR (Copp *et al*., 2014). The library was constructed with an expected frequency of 1.3 nucleotide substitutions per gene distributed across the entire LinB sequence (∼1.5 mutations/kb), corresponding to ≈1 amino acid substitution per protein. With this low mutation frequency, a library was set for single-point variants without positional bias using Poisson sampling at the nucleotide level. Clones carrying frameshifts or premature stop codons were excluded. All plasmids were expressed in the *E. coli* BL21(DE3) strain (New England Biolabs). For overnight growth and protein production in the 96-well microtiter plates, Luria-Bertani (LB) media (Sigma Aldrich) supplemented with a selective antibiotic was used. The expression was induced by IPTG (isopropyl β-d-1-thiogalactopyranoside, Duchefa Biochemie). Chemical cell lysis was performed using SoluLyse® protein extraction reagent (Genlantis), followed by centrifugation. The soluble supernatant was collected and mixed with the GFP_1-10_ fragment in excess. The GFP_1-10_ fragment was produced and purified separately from the inclusion bodies, following the protocol by Cabantous and Waldo (2006) with minor modifications. After 5 h of co-incubation of the GFP fragments and chromophore maturation, fluorescence was measured using a plate reader CLARIOstar® Plus (BMG LABTECH) with excitation/emission settings of 488 and 530 nm, respectively. The individual mutations were determined via Sanger sequencing (PlateSeq, Eurofins Genomics, Germany). Each variant was represented by a single transformant, which was used for three independent protein productions and the split-GFP assays (three independent cultivation replicates).

A significant part of the LinB dataset was collected using a semi-automated robotic screening platform for enzyme libraries (Dörr *et al*., 2016). The system integrates robotic liquid handling, cell cultivation, and downstream analysis in microtiter plates, including a fluorescence readout of the split-GFP assay. This allowed us to apply the same assay principles under more standardized and reproducible conditions, while increasing throughput compared to manual handling (**Figure 2**).

**Figure 2:**
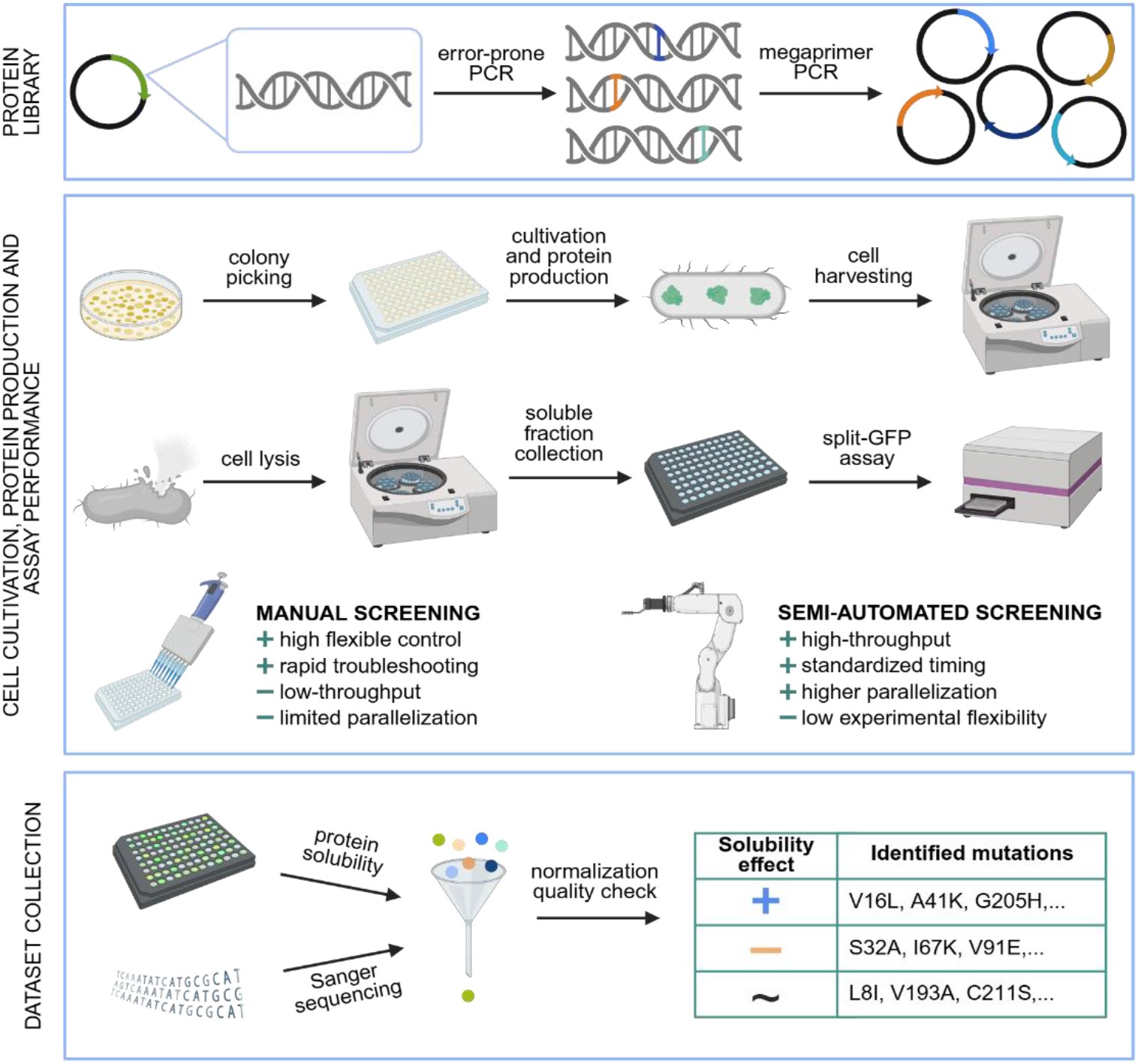
Generation of the LinB mutational solubility dataset. Schematic overview of the experimental workflow used to generate the LinB solubility dataset. **Protein library:** construction of LinB substitution variants fused to the GFP_11_ tag via error-prone PCR and megaprimer PCR. **Cell cultivation, protein production, and assay performance:** expression of genes for protein variants in E. coli, cultivation in the 96-well microtiter plate, production of the haloalkane dehalogenase, and harvesting of the cells via centrifugation, followed by preparation of soluble fractions, and quantification of soluble protein using split-GFP complementation measured as fluorescence. Experiments were performed using both low-throughput in-house microtiter plate screening and high-throughput automated screening on a robotic platform. **Data consolidation:** protein solubility matched with respective mutations identified via Sanger sequencing, followed by normalization, quality control, and compilation of curated mutational solubility datapoints used for downstream analysis and model training.

### Data for external validation

We utilized recently published DMS data for human myoglobin (UniProt: P02144, PDB: 3RGK), obtained via the established YSD assay (Küng *et al*., 2025), and measured in duplicate. Display levels were measured using the same readout class from which the TEM and LGK-YSD training data derive (Table 1), and the caveat that high-throughput scores are proxies that may also track thermodynamic stability applies equally here. The myoglobin set therefore probes soluble expression in the yeast secretory pathway, complementing LinB, which reports cytoplasmic soluble expression in *E. coli*. We created three subsets of the 2351 single-point mutants reported to filter out variants with an ambiguous effect. Based on the mean and standard deviation (SD) of fitness, we kept 34–44% of the datapoints satisfying the following conditions: |*mean*| − *SD* > 0.05 (1037 datapoints), |*mean*| − 3*SD* > 0 (966 data points), and |*mean*| > 0.01 ∧ *SD* < 0.01 (801 datapoints).

### Evaluation protocol

To assess the performance of models, we employed informedness (Powers, 2011) as the primary metric, as it is robust to class imbalance and provides an intuitive scale: 100 % indicates a fully informed predictor, 0 corresponds to a random or naïve predictor, and −100 % denotes a predictor that is always wrong. Informedness can be understood as balanced accuracy (BA) scaled to the range [−100 %, 100 %]. For binary classification, informedness is given by *I*_2_ = *sensitivity* + *specificity* − 1 = 2*BA* − 1 = TP/P + TN/N − 1, where TP and TN are true positives and negatives, P and N are the total positive and negative cases, respectively. In subsets where some class was not represented, and thus informedness was undefined, we used the correct prediction ratio (CPR) as an alternative: *CPR* = #(*correct predictions*) / #(*all predictions*) CPR generalizes accuracy for multiclass classification (Yang *et al*., 2021). We applied bootstrap resampling to each prediction set with 1000 iterations to estimate prediction uncertainty. The resulting 95 % confidence intervals were derived using percentile-based estimation.

## Results

### Cross-evaluation of the models

Our DMS training data consists of three proteins: TEM, LGK, and PKS. We trained models on every combination of these proteins. We also included a model trained solely on the LGK-GFP assay, bringing the total to eight trained models (**Table S1**). Each model was evaluated on all the DMS test subsets (**Figure 3**).

**Figure 3:**
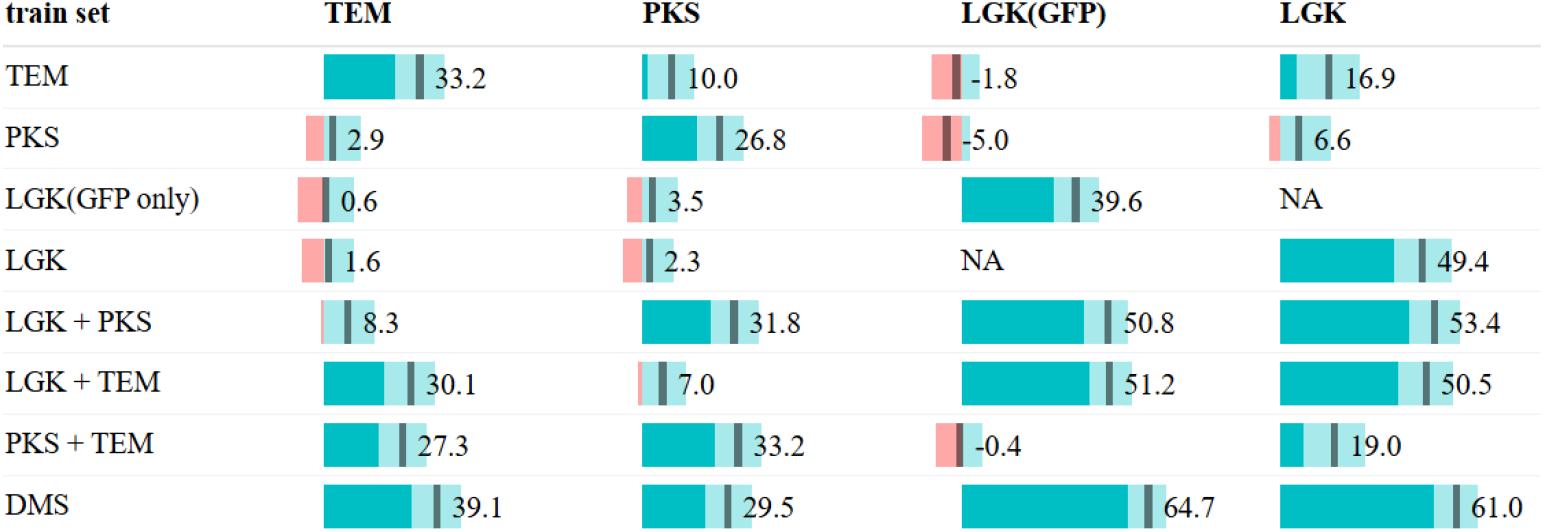
**Cross-evaluation of models** trained on different training subsets of deep mutational scanning (DMS) data. The first group of models is trained on a single protein, the second on two proteins, and the last on all three. No strong generalization was observed across the DMS proteins; the only exceptions were weak but significant generalization to LGK by the TEM (16.9 %) and PKS + TEM (19.0 %) models. The performance metric used is informedness (−100 % to 100 %), with values above and below 0 (better and worse than a random predictor) highlighted in cyan and red, respectively. The lighter cyan signifies 95% confidence intervals.

As can be expected, every model performed best on the protein(s) it was trained on. The single-assay models achieved the following informedness scores on their respective test sets: 26.8 % for PKS, 33.2 % for TEM, and 39.6 % for LGK-GFP. Interestingly, the TEM model outperformed the PKS model, despite having 15 % less data. The DMS model, trained on all three proteins, clearly outperforms the others, achieving informedness scores ranging from 29.5 % for PKS to 64.7 % for the LGK test subsets.

We investigated whether the models could generalize to DMS proteins not included in their training. In most cases, no significant generalization was observed. However, two models, trained on TEM and TEM+PKS, both showed weak generalization to LGK, with informedness of 16.9 % and 19.0 %, respectively. The former also weakly generalized to PKS (10.0 %). While limited in practical significance, these results are statistically significant. Notably, in the case of LGK, generalization was only observed on the curated LGK test set and not on the LGK-GFP data alone.

We also conducted experiments focused on optimizing training parameters and how stricter data splits by mutation position (bypos) affect the performance of our models. Detailed results are provided in the Supplement.

### Synergistic effect of additional training data

To better understand the patterns observed in **Figure 3**, we further examined how including additional proteins or assays improves prediction accuracy for the original target protein (**Figure 4**). Synergistic improvement was observed for all three target proteins, as training on a combination of our three proteins outperforms training on a single protein. In the case of TEM, synergy was observed only when both other proteins were included in the training. Importantly, the synergistic effect was not limited to combining different proteins; it also appears when combining different assays for the same protein, as LGK shows. Despite inconsistencies between the YSD and GFP assays, a model trained on both improves performance from 39.6% to 49.4%.

**Figure 4:**
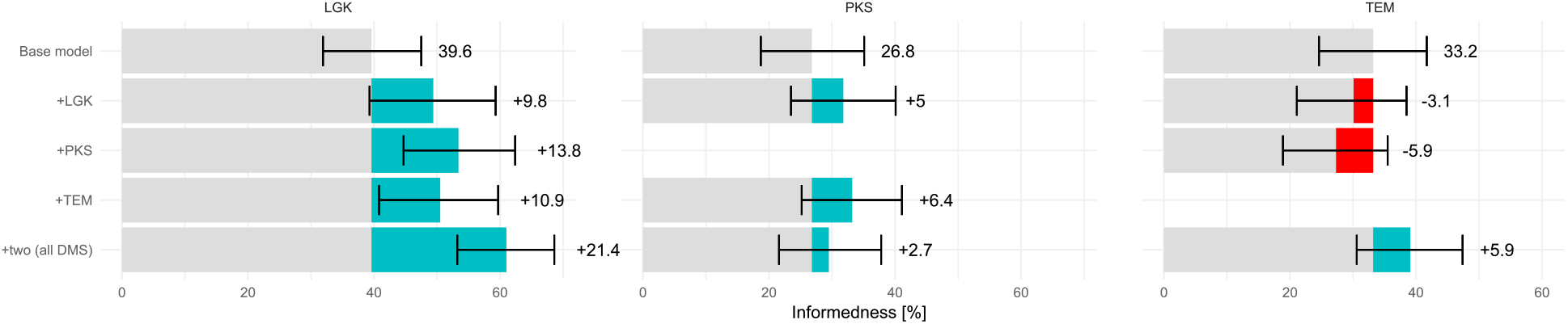
Synergistic effect of additional data inclusion into the training on model performance. The chart shows how the prediction performance of base models, trained for a single protein target, changes when additional protein data are included in the training. Each column represents a specific base model and its corresponding test protein, e.g. the LGK model evaluated on the LGK data. All base models benefited from adding both remaining proteins. When only one was added, the performance of the model on the TEM dataset deteriorated. In the case of the LGK model, adding LGK data refers to incorporating the data from the YSD assay, complementing the GFP data. Error bars are 95 % bootstrap confidence intervals (1000 resamples).

### Independent validation

To assess the generalization capability of our models to proteins unseen in training, we performed a validation on independent data (**Figure 5**). We used the non-DMS test subset, i.e., low-throughput data comprising a diverse set of proteins. This evaluation also included four fine-tuned models: three fully fine-tuned on individual DMS proteins, and one DMS model partially fine-tuned on the non-DMS training data. Only half demonstrated moderate generalization (lower confidence bound greater than 5 %) to the non-DMS proteins. None of them included the TEM data in the final training, and the two worst performers were the models trained for TEM. On the other hand, the best-performing model was the one trained on LGK, with an informedness of 45.6 %. Partial fine-tuning on the non-DMS data was unsuccessful, as the resulting model underperformed even compared to the DMS model from which it was derived.

**Figure 5:**
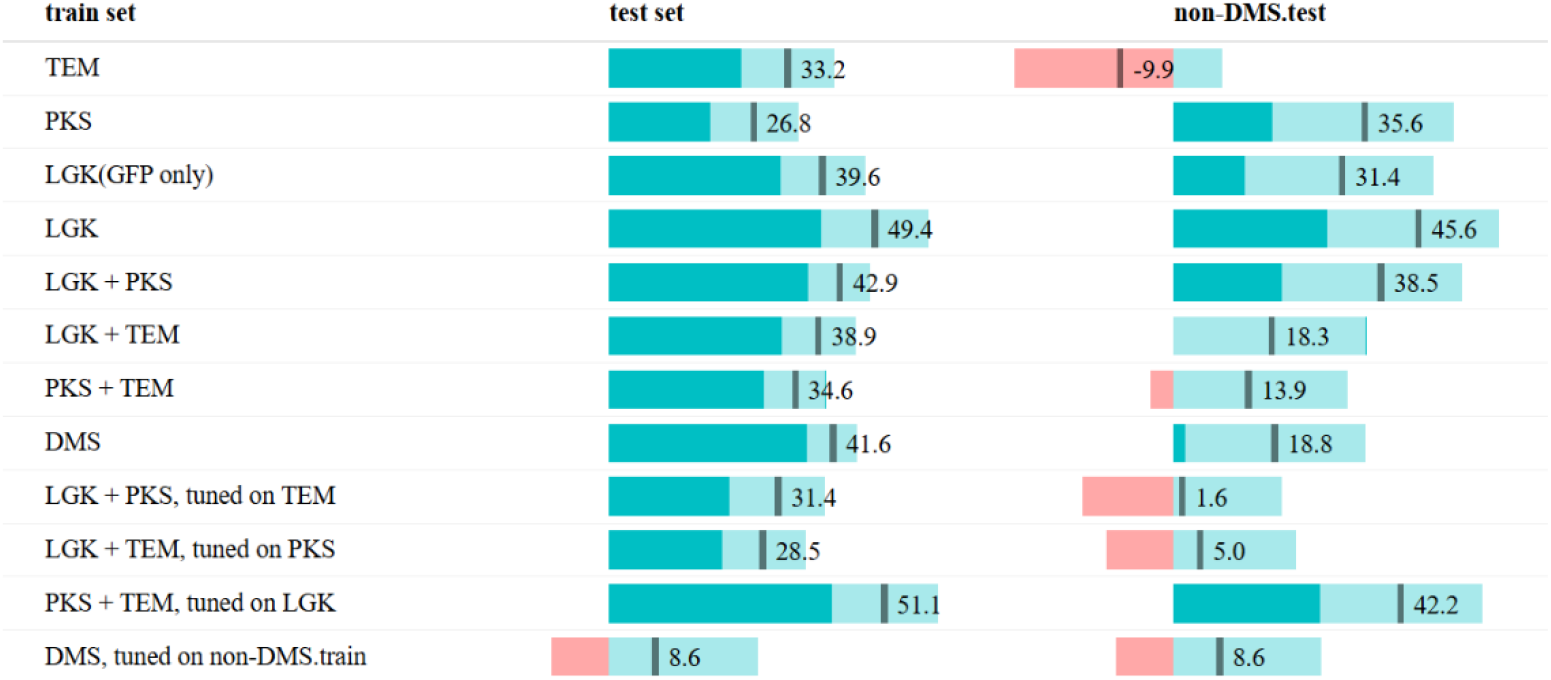
Validation of the trained models. The chart shows the prediction performance of the various trained models on unseen proteins from the test non-DMS data. Some models do demonstrate significant generalization on these never-seen proteins. Interestingly, the models trained with the TEM protein data fail to generalize. The test set in each row corresponds to the results of the respective model on its designated test set, i.e. on the combination of the test subsets of each dataset included in the training, e.g. the TEM test data for the TEM model. Note that the non-DMS fine-tuning data does not overlap with non-DMS proteins from the test set. The performance metric used is informedness (−100 % to 100 %), with values above and below 0 (better and worse than a random predictor) highlighted in cyan and red, respectively. The lighter color signifies 95% confidence intervals.

### Comparison against the state of the art

We compared our models against a selection of existing solubility predictors using unseen data (**Table 2**). We chose two of our models for this evaluation: (a) the LGK-GFP model, the best-performing single-assay model, and (b) the LGK model, the best-performing DMS-based model (**Figure 3**). We included three existing previously published predictors validated for mutational prediction: structure-based OptSolMut (Tian *et al*., 2010) and sequence-based CamSol (Sormanni *et al*., 2015) and PON-Sol2 (Yang *et al*., 2021). The remaining mutational tools, namely PON-Sol (Yang *et al*., 2016), TopLapGBT (Wee *et al*., 2024) and DeepMutSol (Wang *et al*., 2024), were considered but excluded due to unavailability or technical constraints. For completeness, we also included two general-purpose solubility scoring methods: the revised Wilkinson-Harrison method (rWH, Davis *et al*., 1999), an early model for solubility prediction, and SoluProt as a popular representative of absolute solubility predictors. To enable comparison, we derived mutational classifications for rWH and SoluProt (Hon *et al*., 2021) based on the difference in predicted scores between the original and mutant sequences. A complete list of all the tools we considered is provided in **Table S4**.

**Table 2:**
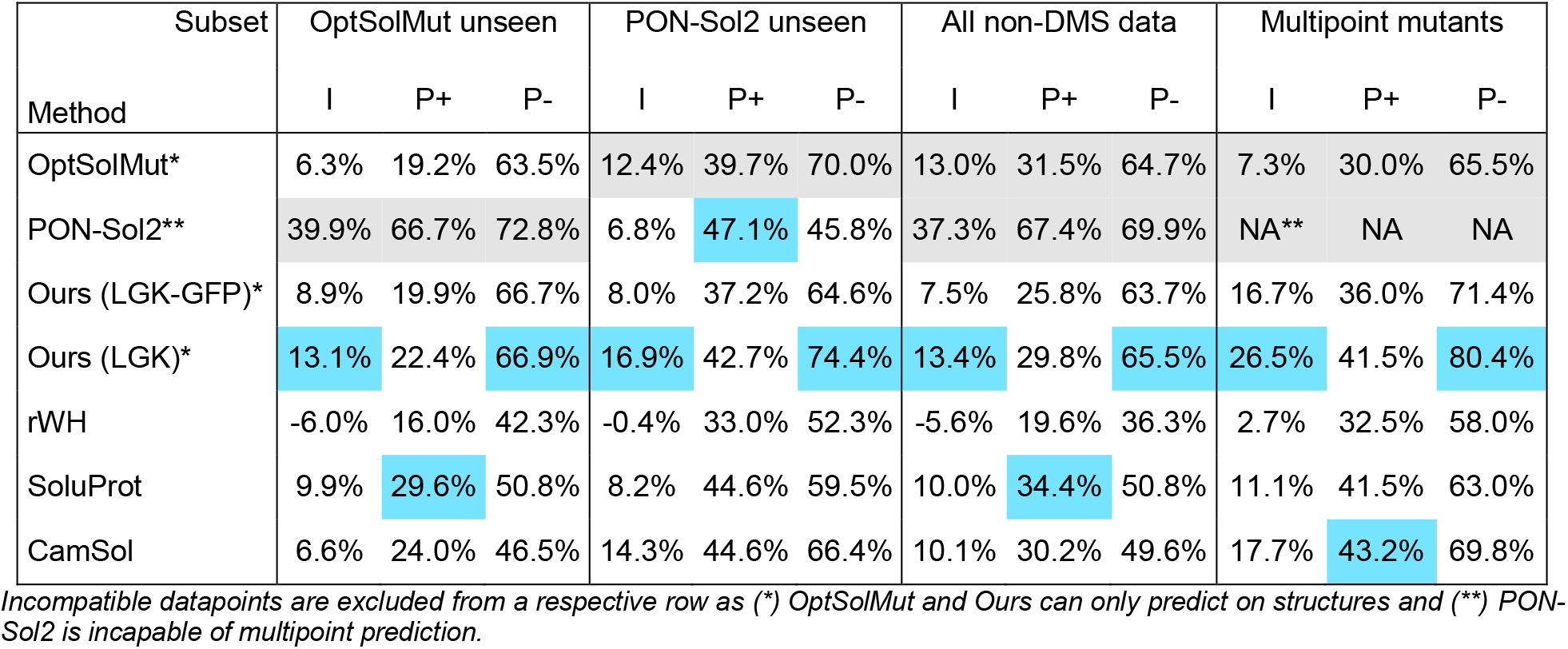
Results of predictors for protein solubility on different subsets of the non-DMS data. The used metrics were informedness (I, ternary, from −100 to 100 %) and precision for the solubilizing/desolubilizing class (P+/P−, 0–100 %). For objective comparison with the OptSolMut and PON-Sol2 predictors, we used subsets that excluded the datapoints seen in the training by the respective tool. When a subset containing such data was used, both tools performed substantially better (as seen in the grey cells). Excluding these, our LGK predictor was leading in overall prediction performance as measured by informedness and precision for the desolubilizing class. Our model showed a considerably stronger performance in the case of multipoint mutants.

| Subset | OptSolMut unseen |  |  | PON-Sol2 unseen |  |  | All non-DMS data |  |  | Multipoint mutants |  |  |
|---|---|---|---|---|---|---|---|---|---|---|---|---|
| Method | I | P+ | P- | I | P+ | P- | I | P+ | P- | I | P+ | P- |
| OptSolMut* | 6.3% | 19.2% | 63.5% | 12.4% | 39.7% | 70.0% | 13.0% | 31.5% | 64.7% | 7.3% | 30.0% | 65.5% |
| PON-Sol2** | 39.9% | 66.7% | 72.8% | 6.8% | 47.1% | 45.8% | 37.3% | 67.4% | 69.9% | NA** | NA | NA |
| Ours (LGK-GFP)* | 8.9% | 19.9% | 66.7% | 8.0% | 37.2% | 64.6% | 7.5% | 25.8% | 63.7% | 16.7% | 36.0% | 71.4% |
| Ours (LGK)* | 13.1% | 22.4% | 66.9% | 16.9% | 42.7% | 74.4% | 13.4% | 29.8% | 65.5% | 26.5% | 41.5% | 80.4% |
| rWH | -6.0% | 16.0% | 42.3% | -0.4% | 33.0% | 52.3% | -5.6% | 19.6% | 36.3% | 2.7% | 32.5% | 58.0% |
| SoluProt | 9.9% | 29.6% | 50.8% | 8.2% | 44.6% | 59.5% | 10.0% | 34.4% | 50.8% | 11.1% | 41.5% | 63.0% |
| CamSol | 6.6% | 24.0% | 46.5% | 14.3% | 44.6% | 66.4% | 10.1% | 30.2% | 49.6% | 17.7% | 43.2% | 69.8% |

*Incompatible datapoints are excluded from a respective row as (\*) OptSolMut and Ours can only predict on structures and (\*\*) PON-Sol2 is incapable of multipoint prediction.*

We used the non-DMS dataset for the comparison, in which datapoints were binned into three classes: desolubilizing, neutral, and solubilizing (−, N, +, respectively). To ensure a fair and objective comparison between predictors, we constructed two evaluation subsets excluding datapoints from either the OptSolMut or the PON-Sol2 datasets. We also created the third subset, which consisted of all the multipoint mutants only. The entire non-DMS dataset was also used for evaluation, as neither of our chosen SoluProtMut models had been trained on these data. Using these four overlapping subsets allowed us to avoid an otherwise excessive reduction of datapoints available for analysis. The performance results for each model across these subsets are in **Table 2**.

Our LGK model outperformed existing solubility predictors in terms of informedness. On the OptSolMut-unseen subset, it exceeded OptSolMut by 6.8 pp; on the PON-Sol2-unseen subset, it exceeded PON-Sol2 by 10.1 pp; and on the full non-DMS dataset, it exceeded SoluProt by 3.4 pp and CamSol by 3.3 pp. Notably, both mutational predictors in the test showed a significant drop in performance when evaluated on unseen proteins, scoring informedness of only 6.8 % and 6.3 %. Surprisingly, SoluProt, a predictor not explicitly designed for mutational analysis, outperformed both, highlighting the limitations of current mutational predictors. Our LGK model, PON-Sol2, SoluProt or CamSol each scored best at least in one of the evaluation categories (highlighted in **Table 2**). The least performing model is almost consistently rWH, often yielding negative informedness. Nonetheless, this is an expected outcome as its simplistic design considers only eight amino acid types. As a result, it misclassifies 161 impactful mutations, treating substitutions among the remaining twelve amino-acid types as neutral.

On the full non-DMS dataset, SoluProtMut gave the highest precision for desolubilizing mutations (65.5 %, against 64.7 % for OptSolMut), with clearer margins on the PON-Sol2-unseen (74.4 %) and multipoint (80.4 %) subsets, whereas SoluProt was the most precise for solubilizing mutations (34.4 %). More importantly, our model was significantly ahead of the competing models in predicting the effects of multipoint mutations, achieving an informedness of 26.5 %, 8.8 pp above the next-best model, CamSol.

### Validation on external data

We collected data on the solubility effect for 277 LinB variants. In addition, we used 1037 mutants from recently published myoglobin data. These datasets were not considered in the development, making them ideal for independent validation (**Figure 6**). We also included PON-Sol2 and CamSol in this validation.

**Figure 6:**
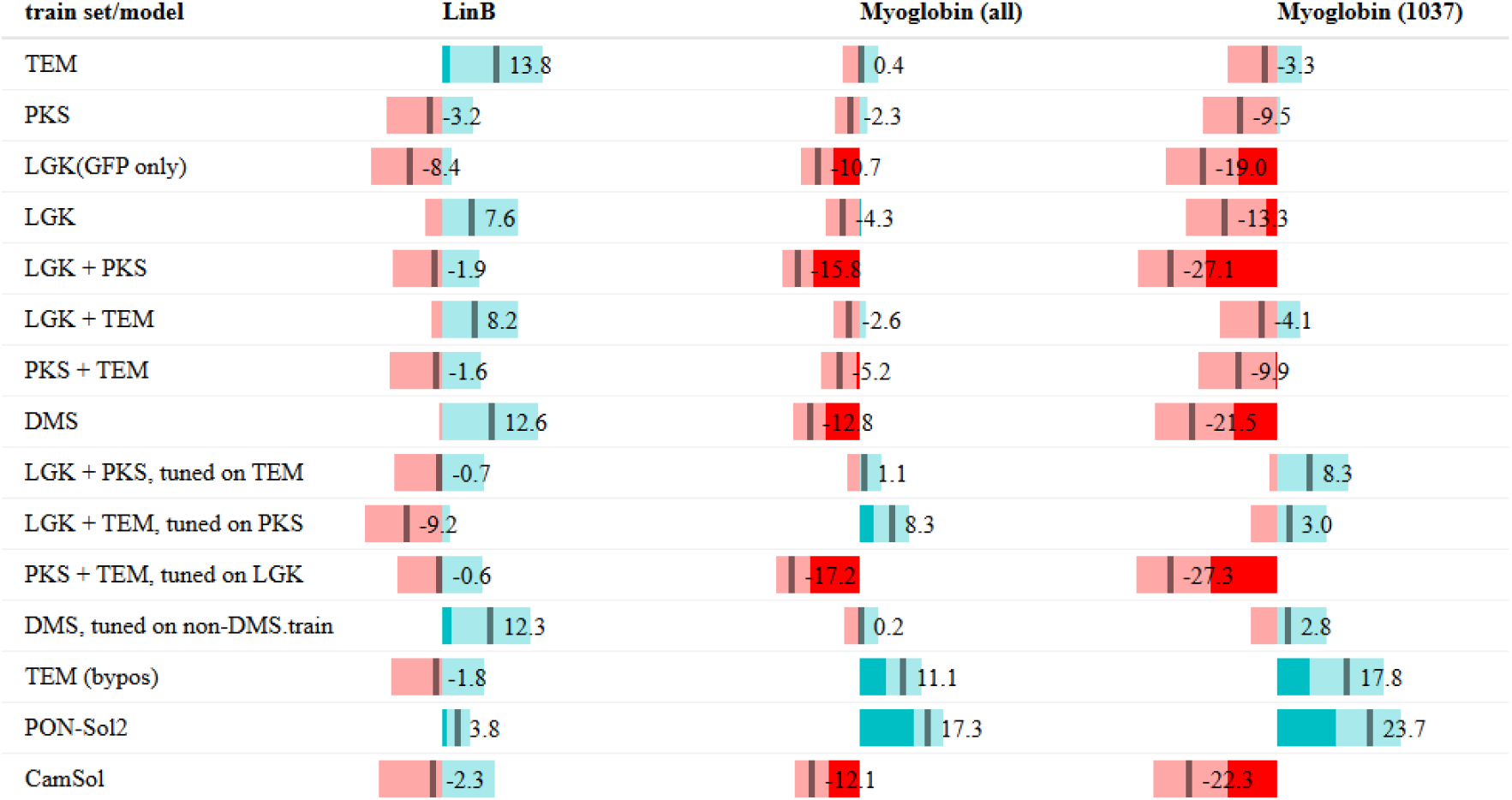
Validation on the external data. (informedness), which comprise mutants of LinB haloalkane dehalogenase and myoglobin. Data for both were collected after the training phase of SoluProtMut. For myoglobin, we show results on the full dataset as well as on a reduced dataset by quality-control filtering. While some of the models showed promising results on the protein-diverse non-DMS data, they struggled to correctly predict mutational effects on these two proteins. The lighter color signifies 95% confidence intervals.

In the case of LinB, the highest informedness was 13.8 % (the TEM model), followed by 12.6 % (the DMS model) and 12.3 % (the fine-tuned DMS model). The remaining models scored below 10 %. In the case of the reduced myoglobin dataset (1037 data points), the highest informedness was 23.7 % for PON-Sol2, followed by 17.8 % for the TEM model with the more stringent data-splitting strategy by position (see the Supplement). Some of the remaining models showed negative informedness, such as the DMS model (−21.5 %) or CamSol (−22.3 %). All but one model showed a larger magnitude of informedness on the reduced dataset than on the full dataset. These conclusions hold for the two alternative reductions as well (**Figure S2**).

### Per-protein accuracy variation

Considering the results on external data, we investigated how the prediction accuracy varies across the 44 proteins of the non-DMS dataset (**Figure 7**). We focused on the LGK model here, as it demonstrated the best performance on the independent test data. After excluding single-example proteins, the model achieved a 100 % success rate on five proteins, and only two proteins showed a 0 % success rate. Our model outperformed random guessing for the majority of the considered non-DMS proteins (23 out of 34). The hardest proteins to predict were those primarily containing mutations classified as neutral, which our binary predictor cannot classify correctly. Both solubilizing and desolubilizing effects were predicted among proteins with multiple mutants. Interestingly, this contrasts with the PON-Sol2 predictions, which generally are unvaried per protein (**Figure S3**).

**Figure 7:**
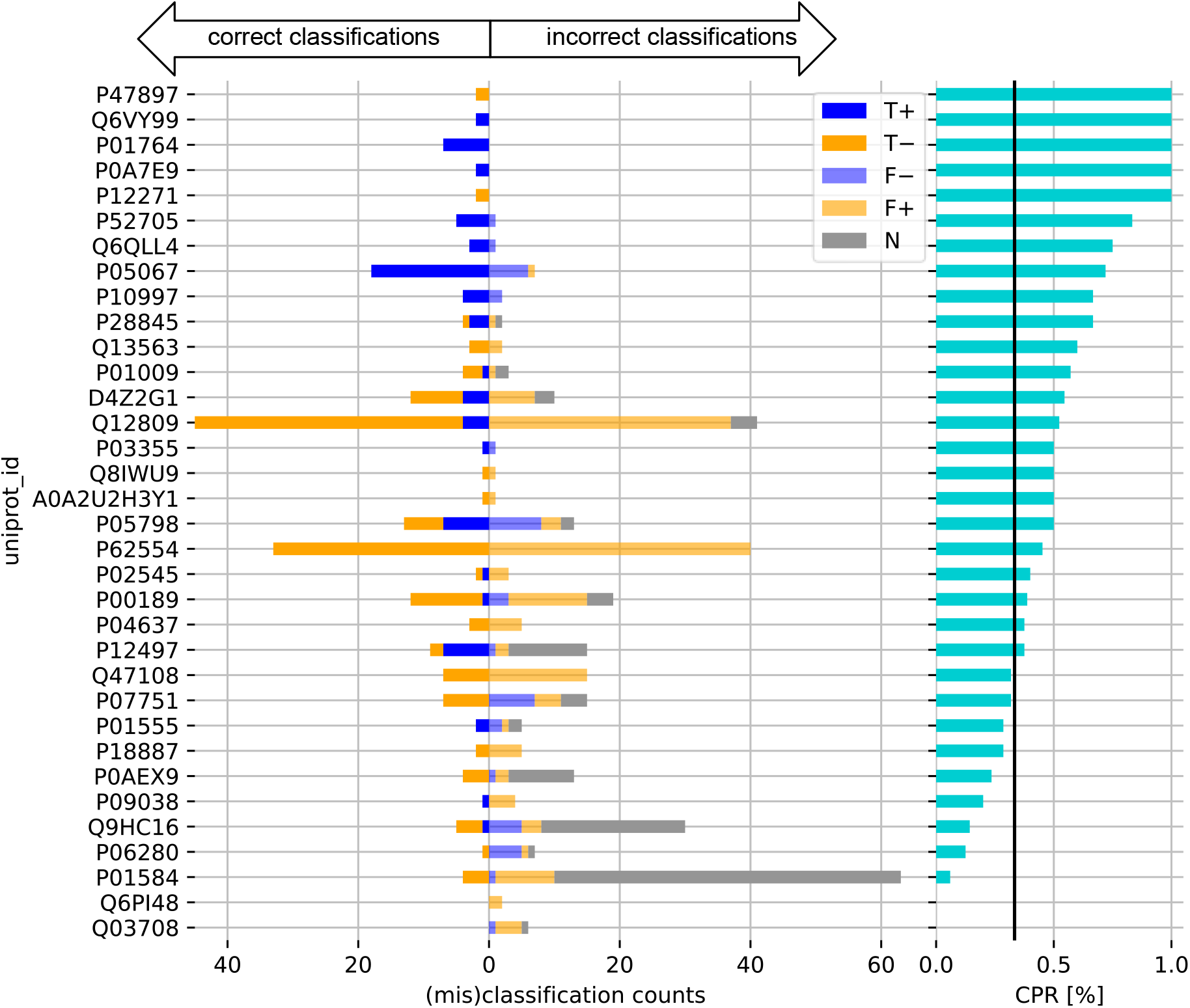
Per-protein performance of our LGK model on unseen proteins. The evaluation on 34 previously unseen non-DMS proteins (n = 553) shows that the predictor performance varies per protein. The left part of the chart shows the per-protein counts of the correctly (saturated colors) and incorrectly (light colors) predicted de/solubilizing (blue/orange) or neutral (gray) effects of mutations. The proteins are sorted by the correct prediction ratio (CPR, the right part) of the model on the given protein. We chose CPR as several proteins contain only a single type of effect. The overall CPR is 44.5 % and our predictor outperforms random guessing for most proteins. For comparison, the CPR corresponding to that of a random predictor (⅓) is drawn by the black line. Note that the proteins with only one example in the dataset were excluded from the chart.

## Discussion

In this work, we leveraged the anti-symmetric design, a property found crucial for the reliability of mutational predictors for stability (Sanavia *et al*., 2020), and recent progress in structure-based deep learning to tackle the problem of protein solubility prediction. Our results show that, on the subsets from which each competitor’s own training data had been removed, SoluProtMut achieved the highest informedness of the methods compared (**Table 2**). The lead of our model is most pronounced in multipoint-mutant effect prediction, where it achieved an informedness of 26.5 %, despite being trained exclusively on single-point mutations.

We substantially extended the available data for mutational solubility prediction by collecting 277 new data points for haloalkane dehalogenase LinB in-house, which we will deposit in SoluProtMut^DB^ (Velecký *et al*., 2022). We used these, along with myoglobin DMS data (Küng *et al*., 2025), for external validation. Additional filtering was needed for the myoglobin data to reduce noise in the validation scores. We tried three criteria, all yielding comparable results (**Figure S2**). Although our models did not generalize well to these data (**Figure 6**), this limitation appears protein-specific rather than systemic, as we observed variation in prediction performance across different proteins (**Figure 7**). LinB was selected as a deliberately demanding validation target: it has a high aggregation propensity and poor solubility at elevated concentrations (Havlásek *et al*., 2025). Importantly, SoluProtMut does not exhibit the per-protein prediction bias, which we noticed with PON-Sol2, that tends to predict a single class per protein (**Figure S3**). This suggests that PON-Sol2 may tend to predict whether a mutated protein is a difficult target for solubility design, rather than the effect of the mutation, although this hypothesis will need further investigation on a larger dataset.

Among the simple models, trained on a single protein, the PKS model showed unexpectedly low test performance, 6.4 pp lower than the TEM model, which has 15 % less training data. This may be attributed to two key differences in the data: lower coverage of the PKS mutational landscape (66 %) compared to TEM (83 %, **Table 1**) and reliance on a homology model rather than an experimentally determined structure. While experimenting with different combinations of the DMS data, we observed that incorporating data from different proteins or even assays not only broadened the scope of a model but also synergistically enhanced model performance (**Figure 4**). A particularly illustrative case is the LGK model, where supplementing the GFP assay data with the additional data from the YSD assay led to notable improvements of +14.2 pp on the independent data. We hypothesize that this is due to an implicit regularization effect from mixing assay data, which introduced data points with inconsistent effects, leading to improved generalization. Strong performance on DMS test subsets does not always translate to generalization. Under the random split, all TEM-based models failed to generalize to non-DMS data (**Figure 5**); the position-wise split recovered generalization for the TEM model (23.7 %, Figure S1). This may stem from the TEM data being derived from the YSD assay, possibly overfitting a model to yeast-specific features. The non-DMS data then predominantly originate from *E. coli*. This explanation is further supported by validation results on myoglobin data, from another YSD assay, in which PON-Sol2 and the TEM (bypos) model significantly outperformed other models (**Figure 6**). Note that a significant part of the PON-Sol2 training dataset is yeast-derived solubility. No similar observation was made with the LGK data, which also contains the yeast measurements.

Our study also highlights the limitations imposed by data scarcity and imbalance. Even though SoluProtMut achieved top-ranking performance in the benchmark experiments, the overall accuracy is moderate, and there is room for improvement, e.g., by increasing the training dataset size (**Figure 4**). Moreover, drawing statistically significant conclusions is difficult due to the wide confidence intervals, sometimes as wide as 38.5 pp for non-DMS data or 30 pp for LinB. The issue of insufficient data on mutational solubility has already been raised in prior studies on predicting solubility (Sormanni *et al*., 2015; Yang *et al*., 2021; Wang *et al*., 2024). Solubility measurements are condition-dependent and influenced by pH, temperature, buffer composition, or an expression system (Baranowski *et al*., 2025). The training data are dominated by measurements obtained in recombinant expression systems, particularly *E. coli*, with a smaller contribution from yeast-based assays. Furthermore, differences in recombinant protein production may arise even between commonly used *E. coli* expression strains due to variations in folding assistance, redox environment, or expression levels. Such effects are not represented explicitly in the current training data and remain an important limitation of existing mutational solubility predictors. As with the previous predictors, our model abstracts away the factors mentioned, thereby avoiding overfitting to rarely annotated experimental conditions. On the other hand, it is a significant limitation of the state of the art, which is probably why no predictor currently performs best across all use cases. For instance, PON-Sol2 or our TEM (bypos) model seems to be a better choice for predicting solubility in yeast over CamSol or our LGK model, which performed best overall. Moreover, even synonymous mutations can affect expression due to codon usage, as manifested by intra-variant score deviations in DMS studies (Klesmith *et al*., 2017; Küng *et al*., 2025), therefore, adding this information to predictors might be another promising strategy to explore.

The model could further benefit from more realistic representations of protein structural variability. In this study, backbone variability is approximated using Gaussian perturbations to coordinates, which provide robustness but do not capture the true conformational landscape in solution. A more faithful description could be achieved using conformational ensembles from molecular dynamics simulations. At the same time, fine-grained effects from local side-chain rearrangements remain beyond the scope of the current architecture. Finally, transfer learning offers another promising path forward. Our network could be pre-trained on related tasks, such as mutational stability prediction, where extensive data are readily available in databases like FireProt^DB^ (Stourac *et al*., 2021) or ProThermDB (Nikam *et al*., 2021). In contrast to solubility data, these databases span hundreds of proteins and provide continuous, high-quality labels, potentially enabling models to learn generalizable protein features through regression, such as protein-folding principles, that also determine solubility. These labels are well-defined quantities of kinetic and thermodynamic stability (Musil *et al*., 2019), providing more homogeneous data. Interestingly, pre-training suits the modular nature of Siamese architecture as it does not require differential data. The base network can therefore also be trained independently on absolute solubility or stability values, leveraging datasets beyond mutation-paired examples.

## Conclusions

This study addresses the challenge of predicting solubility changes upon a mutation. We introduce SoluProtMut, a model based on a novel differential Siamese architecture with implicit anti-symmetry, designed to predict the impact of mutations on solubility using structural data. The underlying convolutional network can learn from 3D protein structures. SoluProtMut is the first model in this domain trained solely on DMS data, yet, on subsets excluding each method’s own training data, it matches or exceeds state-of-the-art methods that were trained, at least in part, on scarce, low-throughput data. Our study therefore shows that high-throughput data are a promising route to overcoming the shortage of mutational solubility data. We believe the new experimental data on the solubility of 277 LinB mutants collected within this study will also help address this problem. However, more comprehensive, less imbalanced solubility data are needed to unlock the full potential of deep learning and enable more statistically robust conclusions. Looking ahead, promising directions include transfer learning from related domains such as protein stability prediction, where abundant data are available. Another interesting direction is incorporating experimental conditions into models, as they strongly influence solubility yet are rarely annotated. Lastly, we encourage future machine learning developers to report confidence intervals when validating their models, as they are essential for interpreting the reliability of predictions.

## Data Availability Statement

SoluProtMut was implemented in Python using PyTorch (Paszke *et al*., 2019), PyTorch Geometric (Fey and Lenssen, 2019) and Biopython (Cock *et al*., 2009) and is available at https://huggingface.co/spaces/loschmidt/SoluProtMut and https://github.com/loschmidt/SoluProtMut. Training of the ensemble of 5 base-network instances takes 1.5 days for the largest data split of all DMS proteins (15 741 training data points) using a single CUDA-enabled GPU (Nvidia 1080 Ti). Inference takes only seconds per mutation on a CPU, i.e., no specialized or high-performance hardware is required. The provided statistics and analyses were created using pandas (McKinney, 2010) and the R reactable, boot, and ggplot2 (Wickham, 2011) libraries. The in-house experimental data are publicly available on Zenodo (10.5281/zenodo.22080131) and will also be released as part of SoluProtMut^DB^ v2.0.

## Acknowledgments

The authors would like to express their thanks to the Czech Ministry of Education, Youth and Sports (TEAMING CZ - CZ.02.01.01/00/23_029/0008437-, EXCELES Onco - LX22NPO5102, ESFRI RECETOX - LM2023069) and the Czech Science Foundation (GX25-17329X). This project has received funding from the European Union’s Horizon 2020 research and innovation programme under grant agreement No. 857560 TEAMING, and by the European Union Centre of Excellence CLARA (101136607), co-funded by the EU from the Operational Programme Jan Amos Komenský (project “Center for Artificial Intelligence and Quantum Computing in System Brain Research", reg. no. CZ.02.01.01/00/23_029/0008437). The article reflects the authors’ view, and the Agency is not responsible for any use that may be made of the information it contains. Computational resources were provided by the e-INFRA CZ and ELIXIR-CZ projects (90254 and LM2023055), supported by the Ministry of Education, Youth and Sports of the Czech Republic. We are grateful to the Deutsche Forschungsgemeinschaft (DFG, INST 292/118-1 FUGG) and the federal state Mecklenburg-Vorpommern for financing the LARA robotic platform. In addition, Brno Ph.D. Talent Scholarship holder Hana Faldynová acknowledges funding from the Brno City Municipality. We thank Vojtěch Voldřich for his contribution to the development of a Python data-processing pipeline for plate reader analysis and the generation of standardized experimental outputs.

## Supplementary Information

**Table S1:** List of training strategies. We constructed 12 strategies for training, of which 8 used different protein-assay subsets of the DMS data and the remaining served for exploring fine-tuning. A strategy (or model) name refers to the name used in the source files.

| *Strategy name* | *Training data* | *Source assays* | *Training objective* |
|---|---|---|---|
| *on-TEM* | TEM | YSD | Single-assay/protein |
| *on-PKS* | PKS | GFP | Single-assay/protein |
| *on-LGK-GFP* | LGK-GFP | GFP | Single-assay/protein |
| *on-LGK-dupl* | LGK | YSD + GFP | Single-protein |
| *on-DMS-leave-LGK* | PKS + TEM | YSD + GFP | Double-protein |
| *on-DMS-leave-PKS* | TEM + LGK | YSD + GFP |  |
| *on-DMS-leave-TEM* | PKS + LGK | YSD + GFP |  |
| *on-DMS* | DMS | YSD + GFP | All-DMS data training |
| *fine-tune-for-LGK* | PKS + TEM, LGK | YSD + GFP | Full-model fine-tuning |
| *fine-tune-for-PKS* | TEM + LGK, PKS | YSD + GFP, GFP |  |
| *fine-tune-for-TEM* | PKS + LGK, TEM | YSD + GFP, YSD |  |
| *fine-tune-DMS* | DMS, non-DMS.train | YSD + GFP, various | Partial-model fine-tuning |

### Hyperparameter optimization

To further enhance the predictive performance of our network, we carried out hyperparameter optimization. The subject of optimization was the on-LGK-GFP training strategy (**Table S1**), which resulted in a 5.3 pp boost in predictive performance. However, this improvement came at the cost of roughly halved generalization (31.4 % to 16.0 %) to the non-DMS data, highlighting a trade-off between overfitting and generalizability (**Figure S1**). The following training and model hyperparameters were systematically explored: weight decay, learning rate, warm-up epoch, feature vector size, global pooling method, and amino acid encoding. We employed a one-parameter-at-a-time (OPAT) strategy in combination with a plateau-based learning rate scheduler to isolate the effects of individual parameters. Among the tested encoding schemes, the VHSE physicochemical descriptor was found to be essential for successful training. Models using one-hot encoding or learned embeddings failed to converge or exhibited poor performance, reaffirming the importance of chemically informed input features.

### Training with position-wise data splits

To investigate the robustness of our network, we evaluated its performance under a more stringent data-splitting strategy. Specifically, we applied a position-wise split to the DMS data, ensuring that each mutated position appeared exclusively in either the training or the test subset, but never in both. This affected model performance as expected. Test performance of the LGK model fell by 31.5 pp to 17.9 %. At the same time, the model generalization to non-DMS data was only marginally affected, decreasing only by 5.4 pp to 40.2 %. In contrast, the PKS and TEM models experienced a collapse in test performance. However, the stricter split unexpectedly recovered generalization for the TEM model: its performance on the non-DMS data increased from −9.9 % to 23.7 % (**Figure S1**).

**Figure S1:**
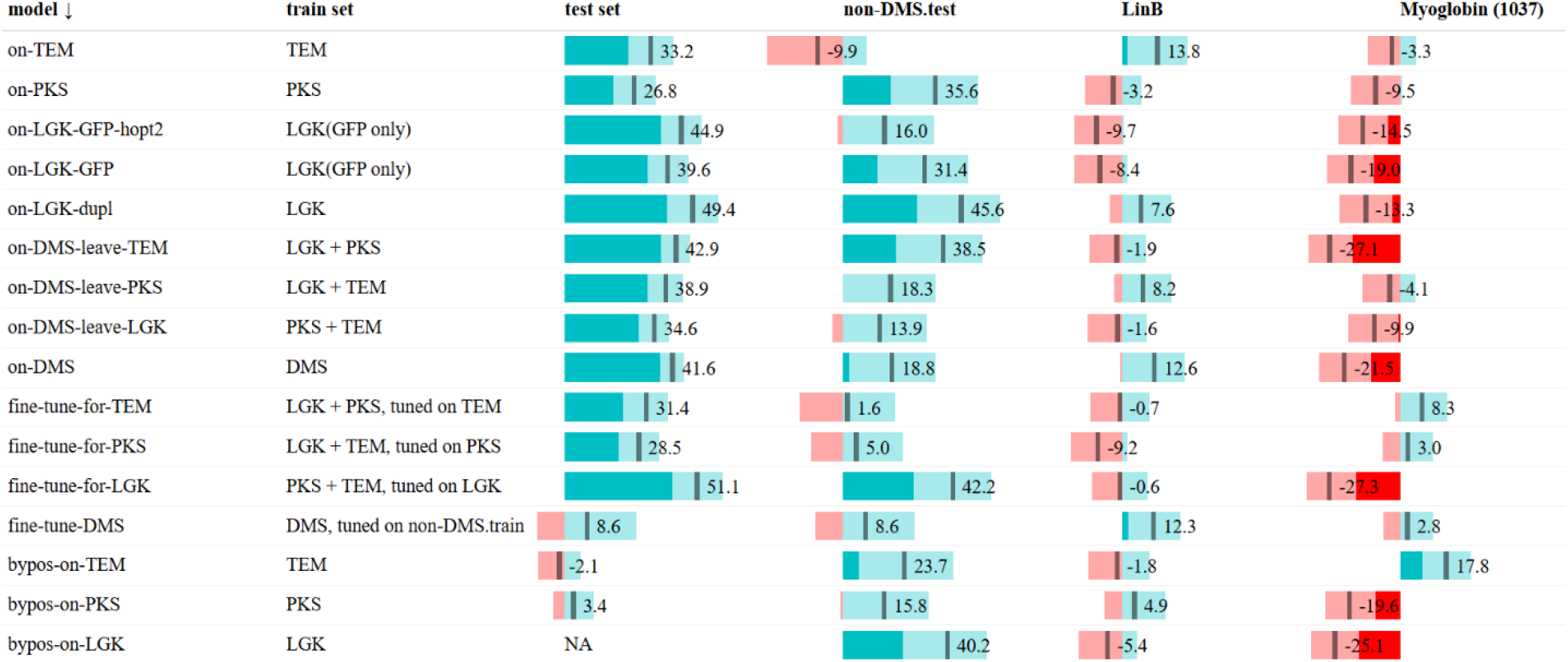
Complete results of all the trained SoluProtMut models. The presented results are the informedness on the designated test set of each model and the three validation sets. The lighter color signifies 95% confidence intervals.

**Figure S2:**
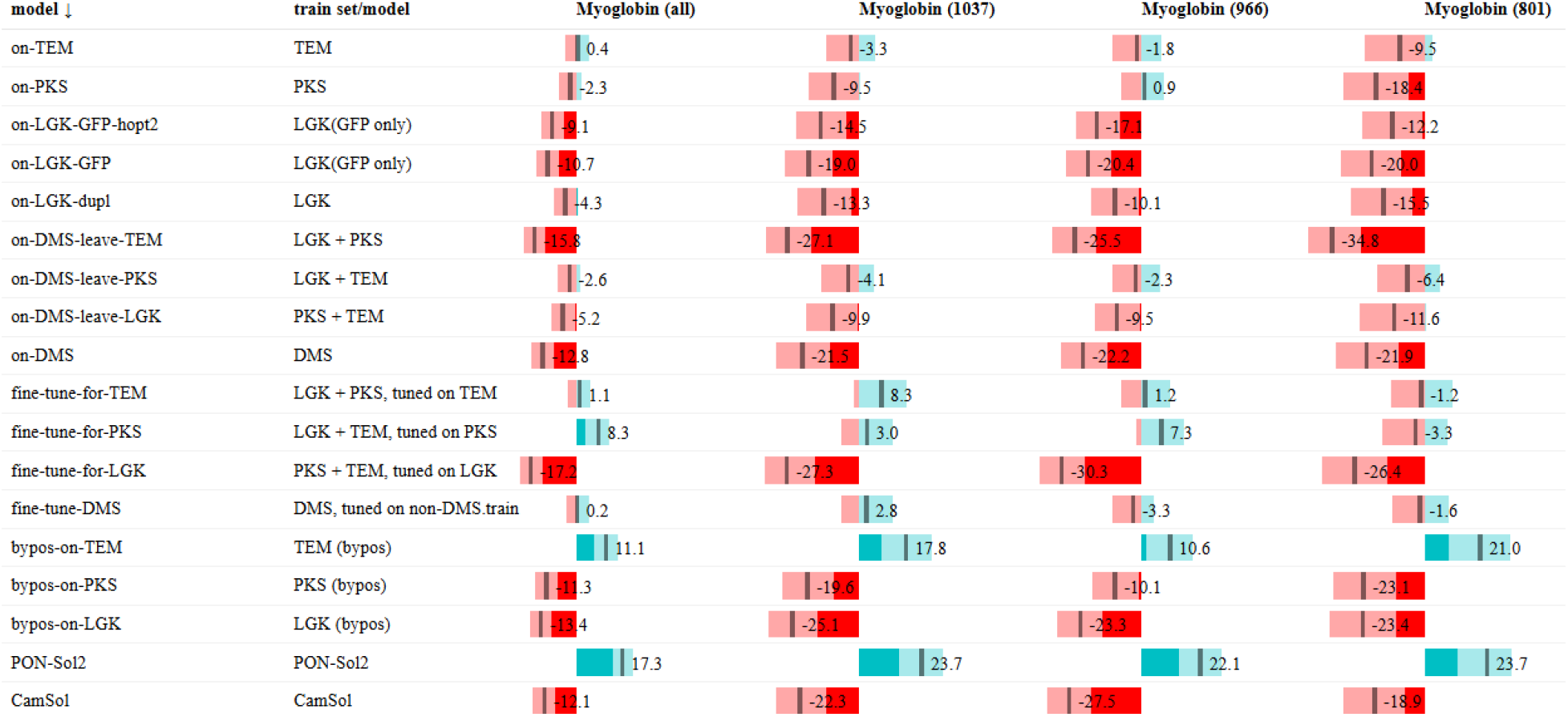
Myoglobin data filtering. All three filtering criteria (down to 1037, 966 or 801 data points) display a higher magnitude of the validation results than the full myoglobin DMS dataset (2351 data points). The lighter color signifies 95% confidence intervals.

**Table S2:** Network design changes introduced in our base network (GCNN). Our network is based on the IEConv network (Hermosilla et al., 2020), which was considerably simplified. AA: amino acid. N/A: not applicable

|  | *IEConv* | *Ours* |
|---|---|---|
| *Protein structure input* | On the atomic level | On the alpha-carbon level |
| *Preprocessing* | Calculation of hydrogen atoms and bonds | N/A |
| *Input node feature representation* | 6D atom feature vector | 8D AA-VHSE embedding |
| *No. of convolution layers* | 2 intra-AA-level layers<br>3 AA-level layers | 3 AA-level layers |
| *Convolution operator* | (Hermosilla *et al.*, 2020) | (Hermosilla *et al.*, 2020) |
| *Convolution parameters* | Euclidean distance<br>Main-chain distance<br>Hydrogen-bond distance | Euclidean distance<br>Main-chain distance |
| *Convolution neurons* | 16/32 | 8 |
| *Convolution distances (Å)* | 3, 6, 8, 12, 16 | 8, 12, 16 |
| *Features per layer* | 64, 128, 256, 512, 1024 | 128, 256, 512 |
| *Sub-AA pooling* | AA-specific average pooling | N/A |
| *AA pooling* | Average pooling, size=2 | Average pooling, size=2 |
| *Ensemble size* | N/A | 5 models |
| *Trainable parameters* | 9.8 million | 2.6 million (×5) |

**Table S3:** Solubility effect classes in SoluProtMut^DB^. Labels, corresponding reported effects on mutant solubility, and the total abundance prior to the filtering.

| *Label* | *Observed effect on solubility* | *# datapoints* |
|---|---|---|
| −− | Highly desolubilizing | 11 388 |
| − | Desolubilizing | 9858 |
| *NE* | Desolubilizing to neutral | 15 |
| *N* | Neutral | 5851 |
| + | Solubilizing | 3931 |
| ++ | Highly solubilizing | 1949 |
|  | Total | 32 992 |

**Figure S3:**
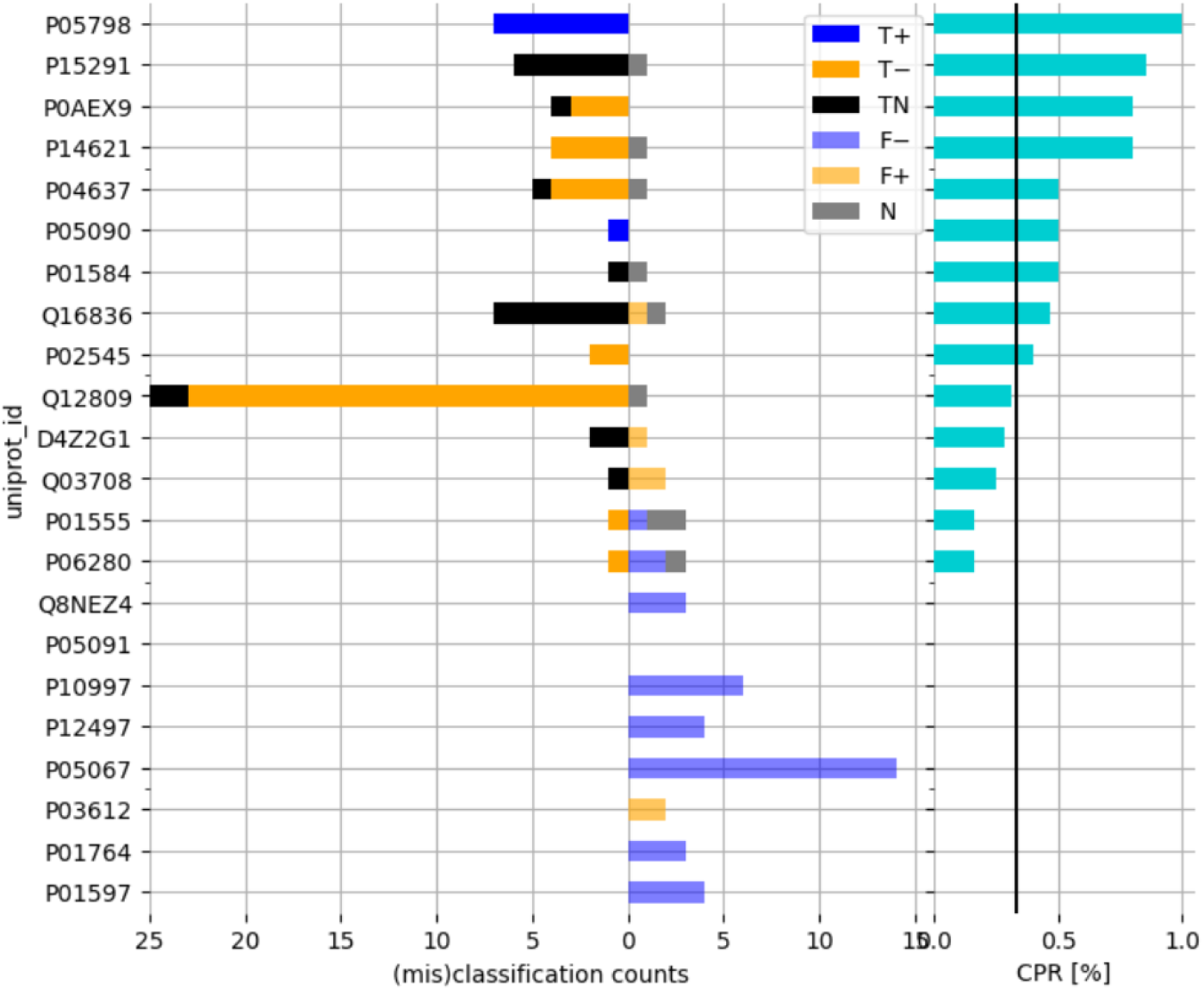
Per-protein performance of PON-Sol2 on unseen proteins. The evaluation was conducted on those non-DMS proteins, previously unseen by PON-Sol2. The left part of the chart shows correct and incorrect classification counts per class. The proteins are sorted by correct prediction ratio (CPR, the right part) of the model on the given protein. The overall CPR is 30.0 %. Notably, PON-Sol2 seems to predict mostly only a single class per protein.

**Table S4:** Predictors of protein solubility considered for comparison with our method. We list known predictors of mutational solubility and two representatives of absolute predictors. PON-Sol, DeepMutSol, and TopLapGBT were not included in our comparison due to unavailability. PON-Sol2, in contrast to tools predicting a continuous score, predicts one of the solubility classes.

| *Method* | *Reference* | *Availability* | *Input* | *Limitation* |
|---|---|---|---|---|
| *Mutational predictors* |  |  |  |  |
| *OptSolMut* | (Tian *et al.*, 2010) | Linux & Mac binary; dataset: archive.math.wsu.edu/math/faculty/bkrishna/DT/OptSolMut/ | PDB |  |
| *PON-Sol* | (Yang *et al.*, 2016) | Tool: unavailable; dataset: structure.bmc.lu.se/VariBench/data/effectspecific/solubility.php | Sequence | Unavailable |
| *PON-Sol2* | (Yang *et al.*, 2021) | Webtool: 139.196.42.166:8010/PON-Sol2/ Python source code; dataset: github.com/XDcat/PON-Sol2 | Sequence | Single-point mutants only; ternary classification: solubilizing, neutral, desolubilizing |
| *DeepMutSol* | (Wang *et al.*, 2024) | Python source code; dataset: github.com/biomed-AI/DeepMutSol | Sequence | Unworkable: trained models and training data not published |
| *TopLapGBT / TopGBT* | (Wee *et al.*, 2024) | Python source code: github.com/ExpectozJJ/TopLapGBT | PDB | Unworkable: insufficient dependency information, hard-coded paths, lacking documentation |
| *Absolute predictors* |  |  |  |  |
| *CamSol* | (Sormanni *et al.*, 2015) | Webtool: www-cohsoftware.ch.cam.ac.uk/ | Sequence, *PDB* | Requires an academic e-mail for registration |
| *SoluProt* | (Hon *et al.*, 2021) | Webtool, source code: loschmidt.chemi.muni.cz/soluprot/ | Sequence |  |

## References

Baranowski, C. et al. (2025) Can protein expression be ‘solved’? Trends in Biotechnology.

Cabantous, S., Terwilliger, T.C., et al. (2005) Protein tagging and detection with engineered self-assembling fragments of green fluorescent protein. Nat Biotechnol, 23, 102–107.

Cabantous, S., Pédelacq, J.-D., et al. (2005) Recent Advances in GFP Folding Reporter and Split-GFP Solubility Reporter Technologies. Application to Improving the Folding and Solubility of Recalcitrant Proteins from Mycobacterium tuberculosis. J Struct Funct Genomics, 6, 113–119.

Cabantous, S. and Waldo, G.S. (2006) In vivo and in vitro protein solubility assays using split GFP. Nat Methods, 3, 845– 854.

Chicco, D. (2021) Siamese Neural Networks: An Overview. In, Cartwright, H. (ed), Artificial Neural Networks, Methods in Molecular Biology. Springer US, New York, NY, pp. 73–94.

Cock, P.J.A. et al. (2009) Biopython: freely available Python tools for computational molecular biology and bioinformatics. Bioinformatics, 25, 1422–1423.

Copp, J.N. et al. (2014) Error-Prone PCR and Effective Generation of Gene Variant Libraries for Directed Evolution. In, Gillam, E.M.J. et al. (eds), Directed Evolution Library Creation: Methods and Protocols. Springer, New York, NY, pp. 3– 22.

Davis, G.D. et al. (1999) New fusion protein systems designed to give soluble expression in Escherichia coli. Biotechnology and Bioengineering, 65, 382–388.

Dörr, M. et al. (2016) Fully automatized high-throughput enzyme library screening using a robotic platform. Biotechnology and Bioengineering, 113, 1421–1432.

Fey, M. and Lenssen, J.E. (2019) Fast Graph Representation Learning with PyTorch Geometric.

Goldenzweig, A. et al. (2016) Automated Structure- and Sequence-Based Design of Proteins for High Bacterial Expression and Stability. Molecular Cell, 63, 337–346.

Havlásek, M. et al. (2025) Decoding Protein Stabilization: Impact on Aggregation, Solubility, and Unfolding Mechanisms. J. Chem. Inf. Model.

He, K. et al. (2016) Deep Residual Learning for Image Recognition. In, 2016 IEEE Conference on Computer Vision and Pattern Recognition (CVPR)., pp. 770–778.

Hermosilla, P. et al. (2020) Intrinsic-Extrinsic Convolution and Pooling for Learning on 3D Protein Structures.

Hon, J. et al. (2021) SoluProt: prediction of soluble protein expression in *Escherichia coli*. Bioinformatics, 37, 23–28.

Klesmith, J.R. et al. (2017) Trade-offs between enzyme fitness and solubility illuminated by deep mutational scanning. PNAS, 114, 2265–2270.

Kramer, R.M. et al. (2012) Toward a Molecular Understanding of Protein Solubility: Increased Negative Surface Charge Correlates with Increased Solubility. Biophysical Journal, 102, 1907–1915.

Küng, C. et al. (2025) Deep mutational scanning reveals a de novo disulfide bond and combinatorial mutations for engineering thermostable myoglobin. Protein Science, 34, e70112.

Magnan, C.N. et al. (2009) SOLpro: accurate sequence-based prediction of protein solubility. Bioinformatics, 25, 2200– 2207.

McKinney, W. (2010) Data Structures for Statistical Computing in Python. Proceedings of the 9th Python in Science Conference, 56–61.

Mei, H. et al. (2005) A new set of amino acid descriptors and its application in peptide QSARs. Peptide Science, 80, 775–786.

Musil, M. et al. (2019) Computational Design of Stable and Soluble Biocatalysts. ACS Catal., 9, 1033–1054.

Nikam, R. et al. (2021) ProThermDB: thermodynamic database for proteins and mutants revisited after 15 years. Nucleic Acids Res, 49, D420–D424.

Paszke, A. et al. (2019) PyTorch: An Imperative Style, High-Performance Deep Learning Library. In, Advances in Neural Information Processing Systems. Curran Associates, Inc.

Powers, D. (2011) Evaluation: From precision, recall and fmeasure to roc, informedness, markedness and correlation. J Mach Learn Tech, 2, 37–63.

Rosace, A. et al. (2023) Automated optimisation of solubility and conformational stability of antibodies and proteins. Nat Commun, 14, 1937.

Sanavia, T. et al. (2020) Limitations and challenges in protein stability prediction upon genome variations: towards future applications in precision medicine. Computational and Structural Biotechnology Journal, 18, 1968–1979.

Shire, S.J. et al. (2004) Challenges in the development of high protein concentration formulations. Journal of Pharmaceutical Sciences, 93, 1390–1402.

Smialowski, P. et al. (2012) PROSO II – a new method for protein solubility prediction. The FEBS Journal, 279, 2192– 2200.

Sormanni, P. et al. (2015) The CamSol Method of Rational Design of Protein Mutants with Enhanced Solubility. Journal of Molecular Biology, 427, 478–490.

Stourac, J. et al. (2021) FireProtDB: database of manually curated protein stability data. Nucleic Acids Research, 49, D319–D324.

Tian, Y. et al. (2010) Scoring function to predict solubility mutagenesis. Algorithms Mol Biol, 5, 33.

Velecký, J. et al. (2022) SoluProtMutDB: A manually curated database of protein solubility changes upon mutations. Computational and Structural Biotechnology Journal, 20, 6339–6347.

Waldo, G.S. et al. (1999) Rapid protein-folding assay using green fluorescent protein. Nat Biotechnol, 17, 691–695.

Wang, J. et al. (2024) Predicting the effects of mutations on protein solubility using graph convolution network and protein language model representation. J Comput Chem, 45, 436–445.

Wee, J. et al. (2024) Integration of persistent Laplacian and pre-trained transformer for protein solubility changes upon mutation. Computers in Biology and Medicine, 169, 107918.

Wickham, H. (2011) ggplot2. WIREs Computational Statistics, 3, 180–185.

Wilkinson, D.L. and Harrison, R.G. (1991) Predicting the solubility of recombinant proteins in Escherichia coli. Biotechnology (N Y), 9, 443–448.

Wrenbeck, E.E. et al. (2019) An Automated Data-Driven Pipeline for Improving Heterologous Enzyme Expression. ACS Synth. Biol., 8, 474–481.

Yang, J. et al. (2015) The I-TASSER Suite: protein structure and function prediction. Nat Methods, 12, 7–8.

Yang, Y. et al. (2016) PON-Sol: prediction of effects of amino acid substitutions on protein solubility. Bioinformatics, 32, 2032–2034.

Yang, Y. et al. (2021) PON-Sol2: Prediction of Effects of Variants on Protein Solubility. International Journal of Molecular Sciences, 22, 8027.

Zhang, Y. et al. (2025) PILOT: Deep Siamese network with hybrid attention improves prediction of mutation impact on protein stability. Neural Networks, 188, 107476.

Zhou, Y. et al. (2023) DDMut: predicting effects of mutations on protein stability using deep learning. Nucleic Acids Research, 51, W122–W128.

